# Dietary betaine-based supplementation improves laying performance and modifies intestinal morphology and microbial profile in hens exposed to a tropical environment

**DOI:** 10.64898/2026.11.13.766587

**Authors:** Zainudin Al Wahid, Adi Ratriyanto, Elham Assadi Soumeh, Sigit Prastowo, Luthfi Adya Pradista

## Abstract

This study evaluated dietary betaine, alone or combined with vitamin C and/or sodium bicarbonate, in laying hens under chronic tropical heat stress, and characterized the intestinal morphological and microbial responses to the best-performing treatment. Two hundred forty 65-week-old Lohmann Brown hens were allotted to five diets (eight replicates of six birds) for seven weeks: basal diet (T0), betaine (T1), betaine + vitamin C (T2), betaine + sodium bicarbonate (T3), and betaine + sodium bicarbonate + vitamin C (T4). The mean temperature-humidity index was 29.68, indicating severe to very severe heat stress. T3 gave the highest hen-day egg production (89.93%) and egg mass (54.83 g), the lowest feed conversion ratio (1.85), and the lowest overall rectal temperature (41.20 vs. 41.57 °C in T0; p < 0.05). T3 was therefore compared with T0 for intestinal histomorphology and microbiota profiling. T3 increased villus height, villus width, and absorptive surface area in the duodenum and jejunum, and villus width and absorptive surface area in the ileum (p < 0.05). Microbiota responses were site-dependent: alpha diversity decreased in the small intestine but was preserved in the cecum, and beta diversity separated T0 from T3 at both sites. *Aeriscardovia aeriphila* increased at both sites, whereas *Romboutsia ilealis*, *Bacteroides togonis*, *Enterococcus cecorum*, and several *Streptococcus* species decreased or became undetectable. Betaine combined with sodium bicarbonate improved laying performance and intestinal morphology and was associated with a marked shift in gut microbial composition under chronic tropical heat stress.

## Introduction

Laying hens in tropical regions still face serious problems due to high ambient temperature and humidity. Prolonged heat exposure disrupts physiological homeostasis and causes heat stress, which negatively affects productivity and health (Salem et al. 2022). In hens, heat stress results in decreased feed intake, hampers the utilization of nutrients, disturbs the acid-base balance, and causes a fall in egg production, all of which lead to substantial economic losses (Farag and Alagawany 2018; Onagbesan et al. 2023). In addition to reducing production, heat stress also affects gut physiology, intestinal structure, and microbial balance (Nawab et al. 2018; Ahmad et al. 2022). As the environmental temperature increases, intestinal blood flow is reduced, the epithelial lining is damaged, and nutrient absorption is impaired, thereby overall digestive efficiency is lowered (He et al. 2018). It also causes a change in composition and metabolic activity of the gut microbiota toward dysbiosis, promoting potentially harmful microorganisms over beneficial ones, which in turn further compromises the gut barrier integrity and nutrient digestion and absorption in the intestine (Nawab et al. 2018; Rychlik 2020).

It is important to have a thorough understanding of the effects of thermal stress on laying hens and develop nutritional strategies to reduce its negative effects. Measures that help to maintain gut function and microbial balance are particularly significant for improving the hens’ ability to cope with heat stress. Betaine (trimethylglycine) acts as both a compatible osmolyte and as a donor of methyl groups. In its role as an osmolyte, betaine assists cells in dealing with osmotic stress by keeping the balance of water inside the cells and by stabilizing protein structure, which in turn lowers the amount of energy needed for cellular maintenance (Ratriyanto and Mosenthin 2018; Ghasemi and Nari 2020; Al Wahid et al. 2024). As a methyl donor, betaine is also involved in one-carbon metabolism and methylation reactions that support protein synthesis, nutrient utilization, and adaptation to environmental stress (Ghasemi and Nari 2020). Dietary betaine supplementation has been shown to enhance laying performance, gut structure, antioxidant status, and heat tolerance in poultry exposed to high ambient temperature (Liu et al. 2019; Attia et al. 2021; Chen et al. 2022).

Sodium bicarbonate (NaHCO_3_) is another nutritional strategy commonly used to counter the physiological effects of heat stress. During heat exposure, birds increase their respiratory rate to dissipate body heat, leading to excessive carbon dioxide loss and potential respiratory alkalosis. Dietary sodium bicarbonate helps restore acid-base and electrolyte balance, thereby supporting physiological stability and performance under hot conditions (Mushtaq et al. 2013; Abbas et al. 2022; Livingston et al. 2022). During panting, CO loss lowers blood pCO and elevates blood pH, causing respiratory alkalosis. The kidneys compensate by excreting bicarbonate (HCO ^-^). Therefore, adding NaHCO does not simply restore acid-base balance by replacing lost bicarbonate. NaHCO is usually used in heat-stressed layers to elevate dietary electrolyte balance (DEB) without adding extra chloride, stimulate water intake for evaporative cooling, and provide carbonate for eggshell formation (Kaya et al. 2004; Yörük et al. 2004; Hassan et al. 2011; Martínez et al. 2021; Abdullah et al. 2025).

Vitamin C exerts antioxidant and anti-inflammatory effects that are particularly valuable during inflammation, oxidative stress, and infection (El-Senousey et al. 2018). Although poultry can synthesize vitamin C endogenously (Whitehead and Keller 2003), high ambient temperature and other stressors increase the physiological requirement for vitamin C while simultaneously reducing its *de novo* synthesis (Van Hieu et al. 2022).

Earlier research demonstrated that betaine, sodium bicarbonate, and vitamin C can improve production responses under heat stress (Al-Qaisi et al. 2023; Kim et al. 2026). However, their effects on intestinal morphology have been inconsistent, and only a limited number of studies have linked these production responses to concurrent changes in the gut microbiota. Furthermore, although the individual effects of betaine, sodium bicarbonate, and vitamin C on heat-stressed poultry are relatively well documented separately, far less is known about how these substances act together. As a result, the combined influence of betaine, vitamin C, and sodium bicarbonate on intestinal structure and microbial balance, especially under prolonged exposure to tropical heat, has not been well investigated. It is also unclear which betaine-based supplementation strategy performs best under tropical field conditions, nor is it known which intestinal and microbial changes accompany the best-performing strategy. Moreover, most studies have used climate-chamber models of acute or cyclic heat exposure, mainly in broilers or in hens at peak lay, whereas evidence from late-phase laying hens kept under naturally occurring, chronic tropical heat is scarce.

The objective of this study was to assess the effects of feeding laying hens a diet supplemented with betaine, either on its own or in combination with vitamin C and sodium bicarbonate, regarding their production performance in tropical environmental conditions. The most effective treatment was then compared with the control group to identify any associated changes in intestinal morphology and gut microbiota composition. It is hypothesized that a dietary supplement containing betaine, either alone or in combination with vitamin C and sodium bicarbonate, would improve production performance, intestinal morphological traits, and the microbial ecosystem of hens exposed to tropical heat, compared with the control.

## Materials and methods

### Ethical approval

The experimental protocol was reviewed and approved by the Ethical Committee for Animal Research of Universitas Sebelas Maret (registration number 18/UN27.20/PT01.01/2023).

### Experimental design and animals

A total of two hundred and forty Lohmann Brown laying hens (65-week-old, average body weight 1.95 ± 0.35 kg) were used in a completely randomized design. The birds were divided into 5 treatment groups, with 8 replicates per treatment and 6 birds per replicate. Pre-treatment adaptation was 1 week, followed by dietary treatments lasting 7 weeks. The hens were housed in a conventional open house battery cage system with 40 cages, each measuring 120 × 60 × 40 cm and holding 6 hens. The chickens were fed once a day, and drinking water was provided ad libitum via nipple drinkers. The basal diet was formulated to meet nutrient requirements for laying hens using breeder’s recommendation (Lohmann Brown-Classic Management Guide; Table 1). The dietary treatments included a basal diet (T0), basal diet supplemented with betaine 1,200 mg/kg (T1), basal diet supplemented with betaine 1,200 mg/kg + vitamin C 500 mg/kg (T2), basal diet supplemented with betaine 1,200 mg/kg + sodium bicarbonate 10 g/kg (T3), and basal diet supplemented with betaine 1,200 mg/kg + sodium bicarbonate 10 g/kg + vitamin C 500 mg/kg (T4).

**Table 1.**
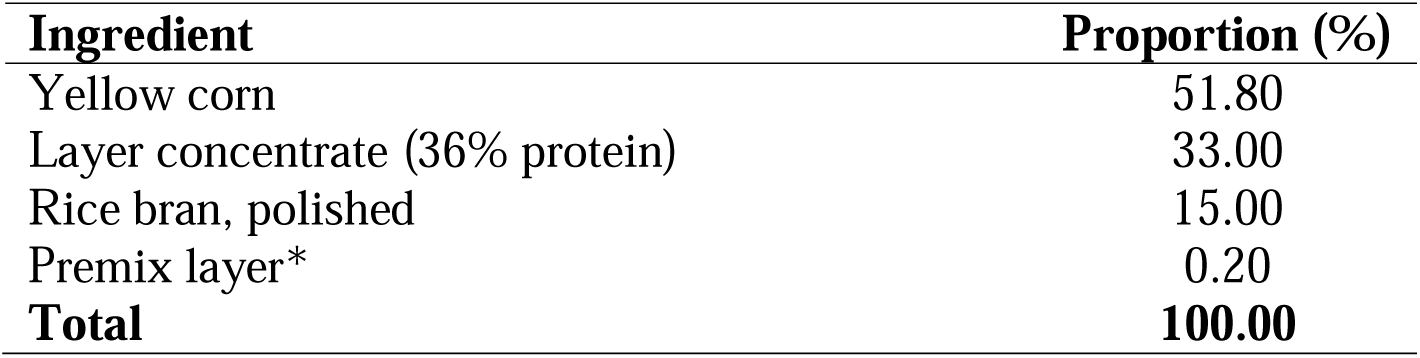
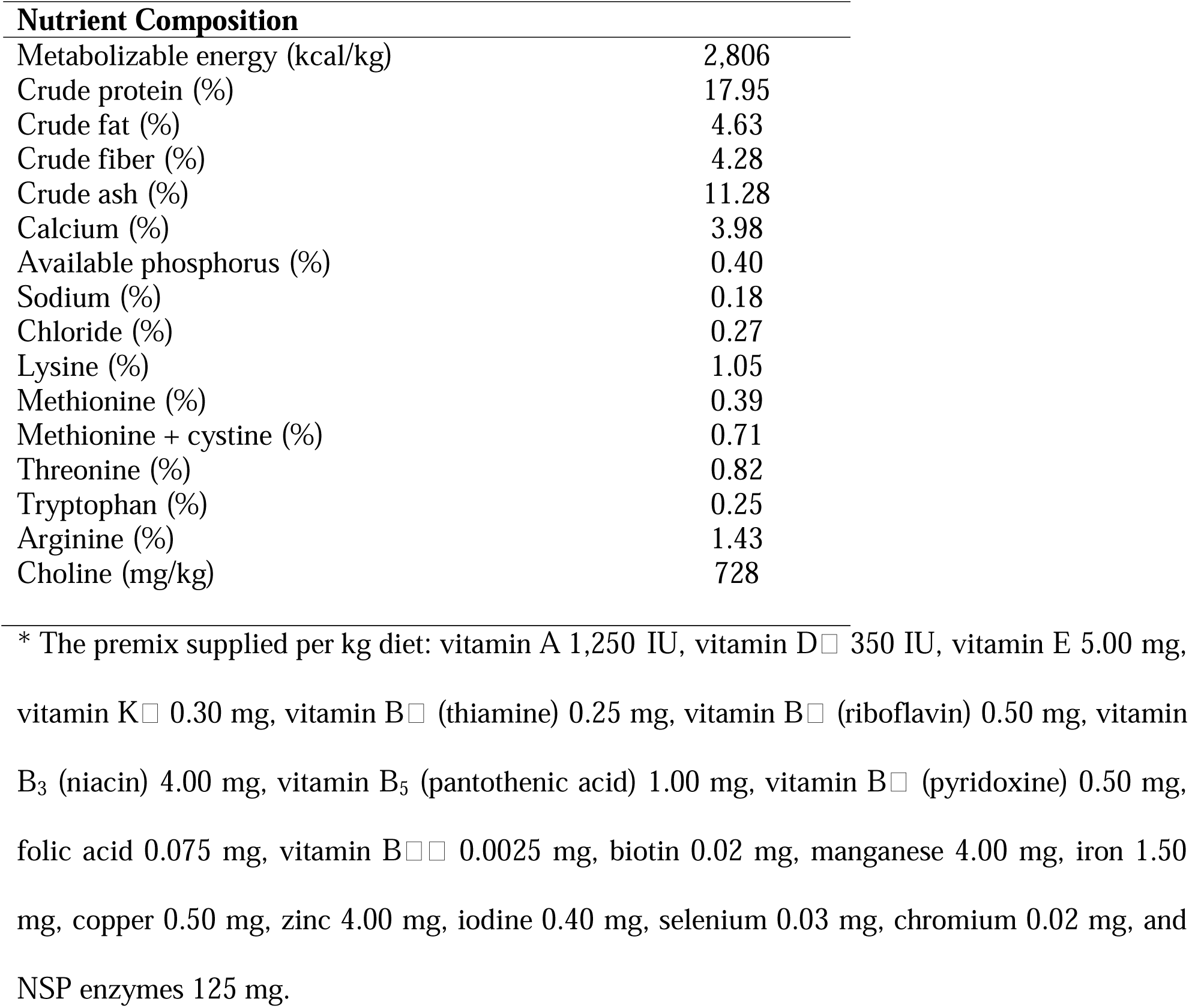
Ingredient and nutritional composition of the basal diet

| Ingredient | Proportion (%) |
|---|---|
| Yellow corn | 51.80 |
| Layer concentrate (36% protein) | 33.00 |
| Rice bran, polished | 15.00 |
| Premix layer* | 0.20 |
| **Total** | **100.00** |
| **Nutrient Composition** |  |
| Metabolizable energy (kcal/kg) | 2,806 |
| Crude protein (%) | 17.95 |
| Crude fat (%) | 4.63 |
| Crude fiber (%) | 4.28 |
| Crude ash (%) | 11.28 |
| Calcium (%) | 3.98 |
| Available phosphorus (%) | 0.40 |
| Sodium (%) | 0.18 |
| Chloride (%) | 0.27 |
| Lysine (%) | 1.05 |
| Methionine (%) | 0.39 |
| Methionine + cystine (%) | 0.71 |
| Threonine (%) | 0.82 |
| Tryptophan (%) | 0.25 |
| Arginine (%) | 1.43 |
| Choline (mg/kg) | 728 |

\* The premix supplied per kg diet: vitamin A 1,250 IU, vitamin D□ 350 IU, vitamin E 5.00 mg, vitamin K□ 0.30 mg, vitamin B□ (thiamine) 0.25 mg, vitamin B□ (riboflavin) 0.50 mg, vitamin B<sub>3</sub> (niacin) 4.00 mg, vitamin B<sub>5</sub> (pantothenic acid) 1.00 mg, vitamin B□ (pyridoxine) 0.50 mg, folic acid 0.075 mg, vitamin B□□ 0.0025 mg, biotin 0.02 mg, manganese 4.00 mg, iron 1.50 mg, copper 0.50 mg, zinc 4.00 mg, iodine 0.40 mg, selenium 0.03 mg, chromium 0.02 mg, and NSP enzymes 125 mg.

### Environmental monitoring

Environmental temperature (°C), relative humidity (%), and wet-bulb temperature were measured continuously using a Lutron data logger (HT-3207SD; Lutron Electronic Enterprise Co., Taipei, Taiwan) placed inside the house at bird level. Data were recorded hourly throughout the study to calculate the temperature-humidity index (THI) and assess potential heat stress. The THI was calculated using the previously reported formula: THI = 0.6 T_db_ + 0.4 T_wb_, where T_db_ = dry-bulb temperature and T_wb_ = wet-bulb temperature (Zulovich 1990).

### Performance data collection

Eggs were collected daily over a period of 7 weeks. The following parameters were recorded: feed intake (FI, g/hens/day), hen-day production (HDP, %), egg weight (EW, g), egg mass (EM, g), and feed conversion ratio (FCR). Hen-day production was calculated as the number of eggs produced divided by the number of hens and then multiplied by 100%. Egg mass was determined by multiplying EW by total hens per treatment, and the FCR was calculated as the ratio of FI to EM (Ratriyanto and Prastowo 2019). Furthermore, rectal temperature was measured weekly with a digital thermometer (Omron MC-343F; Omron Healthcare, Kyoto, Japan) to monitor the birds’ physiological condition.

### Histomorphology analysis

Following the performance trial, intestinal histomorphology was evaluated in the control group (T0) and the best-performing group (T3). Four replicates per group were randomly selected, with one hen per replicate. The purpose of this particular comparison was to determine whether T3’s superior production performance was linked to improvements in the structure of the intestinal mucosa. Because this comparison was selected a posteriori on the basis of production responses, the histomorphological and microbiota results characterize the best-performing treatment rather than the individual effects of each supplement. This method is particularly suitable since the processes involved in tissue processing, sectioning, staining, and morphometric quantification impose practical restrictions on the volume of analysis that can be conducted (Burkholder et al. 2008; He et al. 2018).

Samples were collected from three sections of the small intestine (duodenum, jejunum and ileum) in birds that had been randomly selected from each of the assigned groups. These samples were fixed in 10% neutral buffered formalin, dehydrated by means of a graded series of ethanol, embedded in paraffin, cut into sections, and then stained with hematoxylin and eosin (HE). A cross-section of each sample was obtained at 40× magnification using a digital camera (OptiLab Viewer) connected to the Olympus CX23 microscope. The values for villus height (VH, in µm), villus width (VW, in µm) and crypt depth (CD, in µm) were derived using calibrated imaging software. The villus-to-crypt ratio was calculated by dividing the VH by the mucosal CD (Santos et al. 2019). The absorptive surface area was calculated by applying the following formula: 2π × (VW/2) × VH (Prakatur et al. 2019). Only intact, full-sized villi showing no signs of autolysis or mechanical damage were included in the morphometric measurements to ensure data reliability.

### 16S rRNA gene amplicon sequencing

At the end of the 7-week feeding period, digesta were collected from the control group (T0) and the best-performing group (T3) for microbiota profiling. From each group, 18 hens were randomly selected across all eight replicates. Digesta were collected aseptically from two sites: the small intestine (I), consisting of digesta from the duodenum, jejunum, and ileum, and the cecum (C). For each site, digesta from six hens were pooled in equal amounts to form one composite sample. This gave three composite samples per site per group (3 pools × 6 hens = 18 hens per group) and 12 samples in total (2 groups × 2 sites × 3 pools). Samples were placed in 1.5 mL tubes containing RNA*later* solution (Thermo Fisher Scientific, Waltham, MA, USA) and stored at −20 °C until DNA extraction. The V3–V4 hypervariable region of the 16S rRNA gene was amplified with primers 341F (5′-CCTAYGGGRBGCASCAG-3′) and 806R (5′-GGACTACNNGGGTATCTAAT-3′). Amplicon libraries were prepared from the purified PCR products and sequenced on an Illumina platform to generate 2 × 250 bp paired-end reads (100,000 read pairs per sample) by PT Genetika Science Indonesia (Indonesia). Adapter and primer sequences were removed with Cutadapt v4.6 (minimum read length 150 bp; minimum overlap 10 bp), and read quality was assessed with FastQC and MultiQC. Quality filtering, error correction, merging of paired-end reads, and chimera removal were performed with DADA2 to infer amplicon sequence variants (ASVs). ASVs were classified taxonomically against the SILVA nr99 reference database (release 138.2). Downstream analyses were performed in R (version 4.2.3) using the phyloseq, vegan, MicrobiotaProcess, and microbiomeMarker packages.

Alpha diversity was estimated using the Chao1, Shannon, and Simpson indices. Beta diversity was assessed by principal coordinates analysis (PCoA) based on weighted UniFrac and Bray–Curtis distances, computed separately for each intestinal site. Relative abundances at the species level were expressed as a percentage of the total non-chimeric reads in each sample.

### Statistical analysis

Statistical analyses were performed using RStudio (version 4.5.2; R Core Team 2025). Rectal temperature and production performance data were analyzed by one-way analysis of variance, and differences among treatment means were separated using Duncan’s multiple range test. Intestinal histomorphology, alpha diversity indices, and the relative abundance of individual bacterial species were compared between T0 and T3 within each site using an unpaired Student’s t-test, with the pooled replicate sample as the experimental unit. Differences in microbial community structure between T0 and T3 within each intestinal site were tested using analysis of similarities (ANOSIM), multi-response permutation procedures (MRPP), and permutational multivariate analysis of variance (PERMANOVA) with exact permutations. Because three samples per group were available at each site, only 10 distinct permutations were possible, and the smallest attainable p-value for these tests was 0.100. Differences were considered significant at p < 0.05, and 0.05 ≤ p < 0.10 was considered a tendency.

## Results

### Environmental conditions

During the experiment, the ambient temperature and relative humidity varied; daytime temperatures were frequently above 30°C, and the average THI was 29.68. Figure 1 shows how the ambient temperature, the relative humidity, and the THI varied daily over the experimental period.

**Figure 1.**
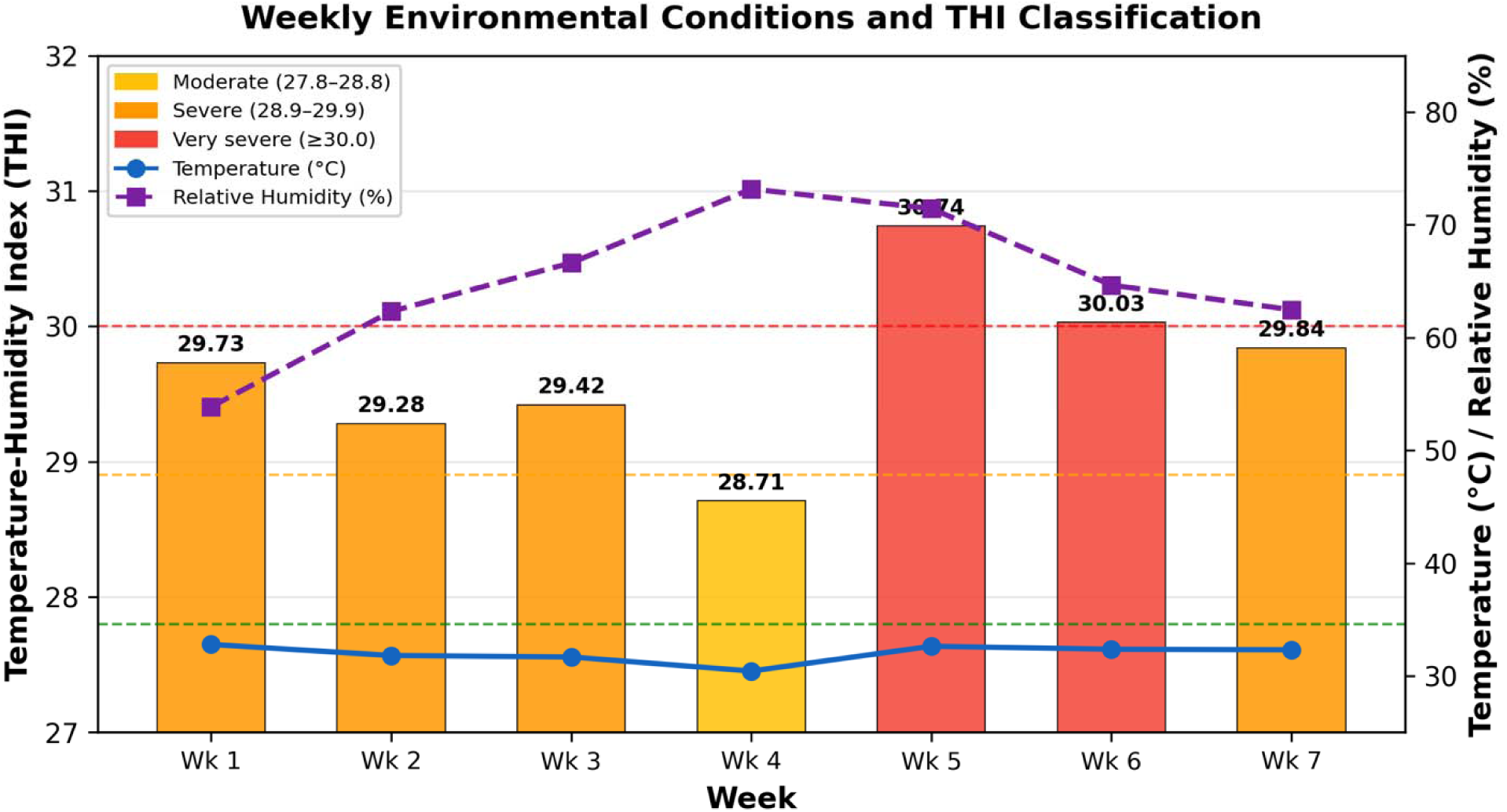
Weekly environmental conditions and temperature-humidity index (THI) classification during the experimental period. Bars represent THI values colored by stress classification: moderate (27.8–28.8), severe (28.9–29.9), and very severe (≥30.0). Lines indicate ambient temperature (°C) and relative humidity (%).

### Rectal temperature

The dietary treatment affected rectal temperature, with differences among treatments becoming significant from week 4 onward (Table 2). During weeks 1–3, rectal temperature tended to differ among treatments (0.05 ≤ p < 0.10; Table 2), with numerically lower values in the supplemented groups than in T0. In week 4, T1, T3, and T4 had lower rectal temperatures than T0 (41.03, 40.90, and 40.97 vs. 41.32°C, respectively; *p* < 0.05), whereas T2 did not differ from T0. In week 5, T3 and T4 had lower rectal temperatures than T0 (41.40 and 41.42 °C, respectively, compared with 41.78 °C; *p* < 0.05), whereas T1 and T2 did not differ from T0. The same pattern was observed in week 6, with T3 and T4 having lower rectal temperatures than T0 (40.88 and 41.07, respectively, compared with 41.32°C; *p* < 0.05). In week 7, T3 and T4 still had lower rectal temperatures than T0 (41.02°C and 41.45°C, respectively, compared to 41.63°C; *p* < 0.05), while T1 and T2 did not differ from T0. Over the entire 7-week period, all supplemented groups had lower rectal temperatures than the control (p < 0.05), with the lowest value in T3 (41.20 °C), which did not differ from T4 (41.31 °C).

**Table 2.** Weekly rectal temperature (°C) of laying hens fed betaine-based supplementation under heat stress conditions

| **Week** | **Temperature (°C)** |  |  |  |  | ***p* value** |
|---|---|---|---|---|---|---|
|  | **T0** | **T1** | **T2** | **T3** | **T4** |  |
| Week 1 | 41.87 | 41.72 | 41.77 | 41.52 | 41.32 | 0.062 |
| Week 2 | 41.67 | 41.43 | 41.65 | 41.33 | 41.52 | 0.092 |
| Week 3 | 41.38 | 41.25 | 41.60 | 41.31 | 41.42 | 0.069 |
| Week 4 | 41.32<sup>a</sup> | 41.03□<sup>c</sup> | 41.17<sup>ab</sup> | 40.90□ | 40.97□<sup>c</sup> | 0.005 |
| Week 5 | 41.78□ | 41.58<sup>ab</sup> | 41.62<sup>ab</sup> | 41.40<sup>b</sup> | 41.42<sup>b</sup> | 0.032 |
| Week 6 | 41.32<sup>a</sup> | 41.10<sup>ab</sup> | 41.15<sup>ab</sup> | 40.88<sup>b</sup> | 41.07□ | 0.009 |
| Week 7 | 41.63□ | 41.58□ | 41.67□ | 41.02□ | 41.45□ | <0.001 |
| Week 1-7 | 41.57<sup>a</sup> | 41.40<sup>b</sup> | 41.51<sup>b</sup> | 41.20<sup>c</sup> | 41.31<sup>bc</sup> | <0.001 |

<sup>a,b,c</sup> Means within a column with different superscript letters differ significantly (p < 0.05).
T0: basal diet; T1: basal diet supplemented with betaine 1,200 mg/kg; T2: basal diet supplemented with betaine 1,200 mg/kg + vitamin C 500 mg/kg; T3: basal diet supplemented with betaine 1,200 mg/kg + sodium bicarbonate 10 g/kg; T4: basal diet supplemented with betaine 1,200 mg/kg + sodium bicarbonate 10 g/kg + vitamin C 500 mg/kg.

### Production performance

Dietary treatment significantly affected FI, HDP, EW, EM, and FCR (*p* < 0.05; Table 3). Feed intake was highest in the control (T0, 109.82 g/hen/day) and lowest in T1 (97.82 g), with all supplemented groups consuming less feed than T0 (*p* < 0.05). HDP was highest in T3 (89.93%) and T4 (88.24%), and lowest in the control (85.11%). Egg weight was higher in T4 than in T0 (*p* < 0.05), whereas T3 did not differ from T0, and both T1 and T2 produced lighter eggs than T0, albeit of small magnitude. Egg mass followed a pattern similar to HDP: T3 and T4 had the highest egg mass (54.83 ± 0.43 and 54.30 ± 0.39 g, respectively), followed by T0 and T2 (52.84 ± 0.45 and 52.57 ± 0.45 g), while T1 recorded the lowest value (50.64 ± 0.44 g). All supplemented groups had a lower FCR than the control (2.09; *p* < 0.05), with T3 achieving the best FCR (1.85).

**Table 3.** Production performance of laying hens fed betaine-based supplementation under heat stress conditions

| **Variable** | **T0** | **T1** | **T2** | **T3** | **T4** | ***p* value** |
|---|---|---|---|---|---|---|
| Feed Intake (g/hen/day) | 109.82 ± 6.11<sup>a</sup> | 97.82 ± 8.13<sup>d</sup> | 101.88 ± 7.81<sup>c</sup> | 100.86 ± 8.57<sup>cd</sup> | 104.97 ± 6.97<sup>b</sup> | < 0.01 |
| HDP (%) | 85.11 ± 5.40<sup>c</sup> | 86.89 ± 6.26<sup>bc</sup> | 86.98 ± 5.32<sup>bc</sup> | 89.93 ± 4.35<sup>a</sup> | 88.24 ± 4.09<sup>ab</sup> | < 0.01 |
| Egg weight (g) | 60.90 ± 1.00<sup>b</sup> | 59.60 ± 1.08<sup>d</sup> | 60.46 ± 0.95<sup>c</sup> | 60.95 ± 0.79<sup>b</sup> | 61.54 ± 0.81<sup>a</sup> | < 0.01 |
| Egg mass (g) | 52.84 ± 0.45□ | 50.64 ± 0.44□ | 52.57 ± 0.45□ | 54.83 ± 0.43□ | 54.30 ± 0.39□ | < 0.01 |
| FCR | 2.09 ± 0.17<sup>a</sup> | 1.99 ± 0.19<sup>b</sup> | 1.94 ± 0.17<sup>b</sup> | 1.85 ± 0.15<sup>c</sup> | 1.93 ± 0.16<sup>b</sup> | < 0.01 |

<sup>a,b,c,d</sup> Means within a column with different superscript letters differ significantly (*p* < 0.05).
HDP: hen-day production; FCR: feed conversion ratio; T0: basal diet; T1: basal diet supplemented with betaine 1,200 mg/kg; T2: basal diet supplemented with betaine 1,200 mg/kg + vitamin C 500 mg/kg; T3: basal diet supplemented with betaine 1,200 mg/kg + sodium bicarbonate 10 g/kg; T4: basal diet supplemented with betaine 1,200 mg/kg + sodium bicarbonate 10 g/kg + vitamin C 500 mg/kg.

### Intestinal histomorphology

Dietary supplementation with betaine and sodium bicarbonate (T3) significantly altered intestinal morphology relative to the control (T0) across all 3 segments of the small intestine (Table 4; *p* < 0.05). In the duodenum and jejunum, T3 increased VH, VW, CD, and absorptive surface area (*p* < 0.05), whereas the villus-to-crypt ratio did not differ significantly between treatments in these segments (*p* > 0.05). In the ileum, T3 increased VW and absorptive surface area (*p* < 0.001), while VH, CD, and the villus-to-crypt ratio did not differ from the control.

**Table 4.** Intestinal histomorphology measurements of laying hens in T0 (control) and T3 (betaine + sodium bicarbonate)

| Intestinal Segment | Variable | T0 | T3 | *p* value |
|---|---|---|---|---|
| Duodenum | Villus height (µm) | 2,158.25 ± 195.82 | 3,030.00 ± 75.14 | < 0.001 |
|  | Villus width (µm) | 334.75 ± 57.47 | 761.25 ± 66.30 | < 0.001 |
|  | Crypt depth (µm) | 355.50 ± 39.08 | 474.25 ± 38.28 | 0.005 |
|  | Villus/crypt ratio | 6.11 ± 0.64 | 6.41 ± 0.38 | 0.444 |
|  | Absorptive surface area (mm<sup>2</sup>) | 2.26 ± 0.35 | 7.26 ± 0.79 | < 0.001 |
| Jejunum | Villus height (µm) | 1,143.50 ± 110.44 | 1,962.00 ± 166.08 | < 0.001 |
|  | Villus width (µm) | 187.25 ± 11.32 | 492.75 ± 35.37 | < 0.001 |
|  | Crypt depth (µm) | 187.75 ± 77.42 | 386.75 ± 81.58 | 0.012 |
|  | Villus/crypt ratio | 7.14 ± 3.64 | 5.22 ± 1.00 | 0.348 |
|  | Absorptive surface area (mm<sup>2</sup>) | 0.67 ± 0.09 | 3.03 ± 0.29 | < 0.001 |
| Ileum | Villus height (µm) | 1,204.00 ± 62.09 | 1,217.75 ± 15.52 | 0.682 |
|  | Villus width (µm) | 239.25 ± 20.76 | 462.50 ± 28.10 | < 0.001 |
|  | Crypt depth (µm) | 194.25 ± 44.71 | 228.25 ± 12.55 | 0.193 |
|  | Villus/crypt ratio | 6.43 ± 1.44 | 5.35 ± 0.32 | 0.192 |
|  | Absorptive surface area (mm<sup>2</sup>) | 0.91 ± 0.11 | 1.77 ± 0.09 | < 0.001 |

T0: basal diet; T3: basal diet supplemented with betaine 1,200 mg/kg + sodium bicarbonate 10 g/kg.

### Intestinal microbiota composition

#### Alpha diversity

Alpha diversity of the intestinal and cecal microbiota was assessed using the Chao1, Shannon, and Simpson indices (Figure 2). In the intestine, all three indices were consistently lower in T3 than in T0 (*p* <0.05). The Chao1 index declined from 190–210 in the T0 intestine to 120–140 in the T3 intestine, the Shannon index decreased from 3.95–4.05 (T0) to approximately 3.52–3.59 (T3), and the Simpson index declined from 0.955–0.965 (T0) to 0.935–0.945 (T3). In the cecum, by contrast, the Chao1, Shannon, and Simpson indices were similar between treatments, remaining high in both T0 (Chao1 ≈ 530–560; Shannon ≈ 5.45–5.50; Simpson ≈ 0.99) and T3 (Chao1 ≈ 540–570; Shannon ≈ 5.45–5.50; Simpson ≈ 0.99).

**Figure 2.**
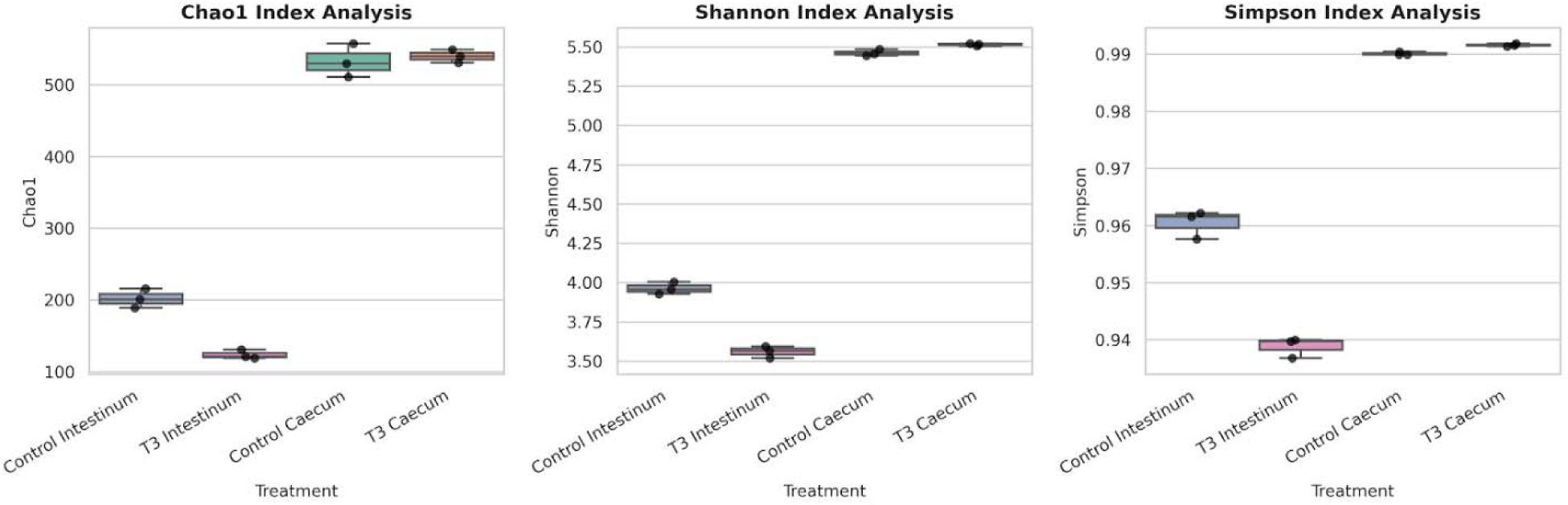
Alpha diversity indices (Chao1, Shannon, and Simpson) of gut microbiota in the intestine and cecum of control (T0) and treatment (T3) groups. Box plots display median, interquartile range, and individual data points.

#### Beta diversity

Principal coordinates analysis (PCoA) based on weighted UniFrac distances revealed distinct clustering of samples by intestinal site and treatment group (Figure 3). Samples from the intestine and cecum formed separate clusters in the PCoA plots. In the small intestine, control and supplemented groups separated completely along PCoA1 (61.31% of variance), forming distinct, non-overlapping clusters (Figure 3, top). In the cecum, the primary axis pair—PCoA1 and PCoA2, accounting for 61.31% and 19.08% of variance, respectively—showed partially overlapping spatial distributions between the control and betaine + sodium bicarbonate groups. However, separation between the cecal control and supplemented groups was fully resolved along PCoA3 (4.61% of variance), with the two groups forming non-overlapping clusters (Figure 3, bottom).

**Figure 3.**
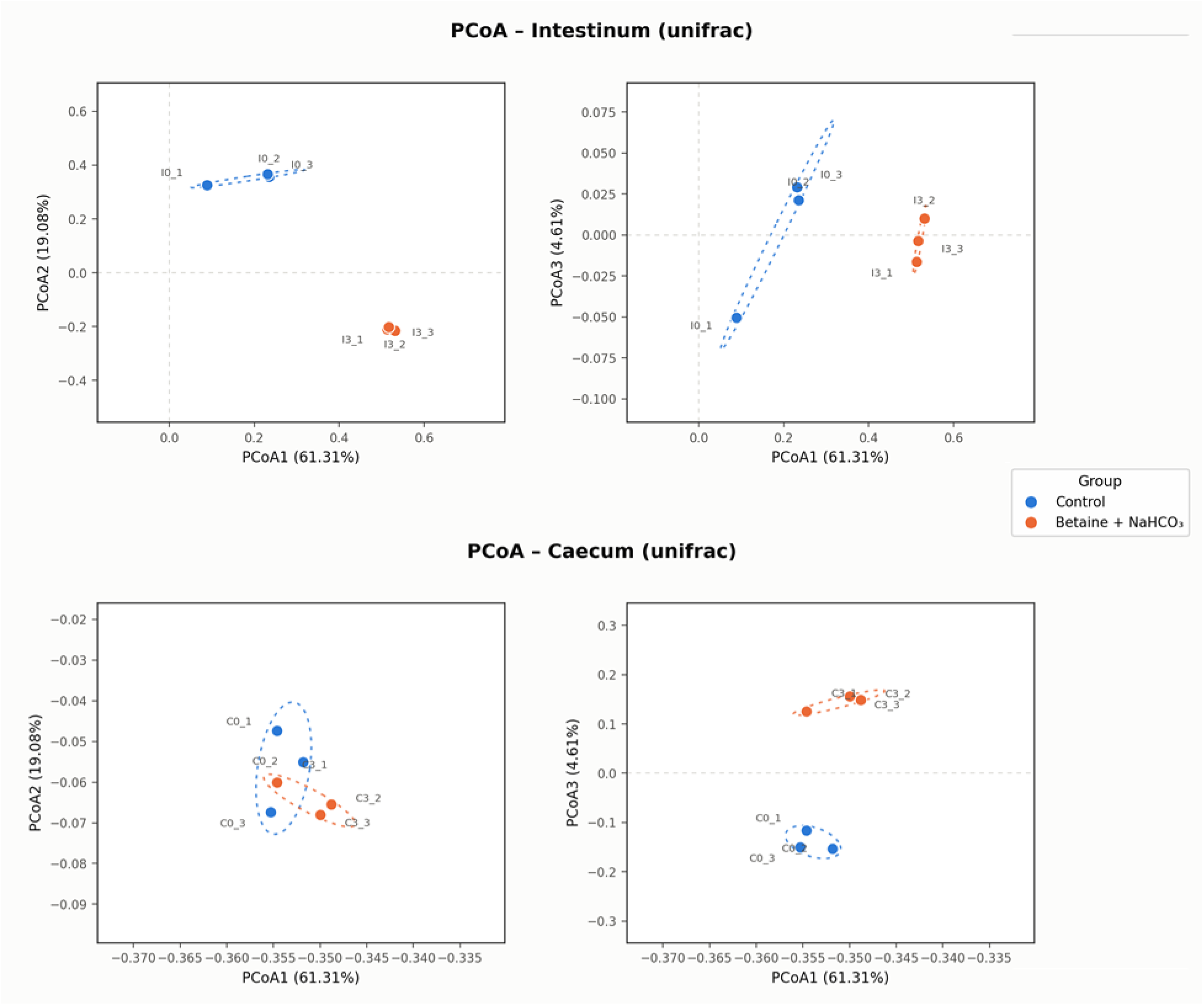
Principal coordinate analysis (PCoA) of weighted UniFrac distances partitioned by tissue site. Top: Small intestine; Bottom: Cecum. Left panels depict PCoA1 vs. PCoA2; right panels depict PCoA1 vs. PCoA3. Blue circles represent control samples; orange circles represent betaine + sodium bicarbonate combination treatment (n = 3 per group). Dashed ellipses represent group dispersion (mean ± 1.6 SD). Axis labels denote the percentage of total variance explained.

To statistically evaluate group separation, Analysis of Similarities (ANOSIM), Multi-Response Permutation Procedures (MRPP), and Permutational Multivariate Analysis of Variance (PERMANOVA via exact permutation) were calculated within each intestinal segment. All three tests yielded the maximum separation statistics attainable with this design (ANOSIM R = 1.000 at both sites; Table 5); however, with three samples per group, the exact permutation p-value could not fall below 0.100, so these differences are descriptive rather than statistically confirmed. PERMANOVA pseudo-*F* values were over four-fold higher in the small intestine (*F* = 78.46) than in the cecum (*F* = 19.46).

**Table 5.**
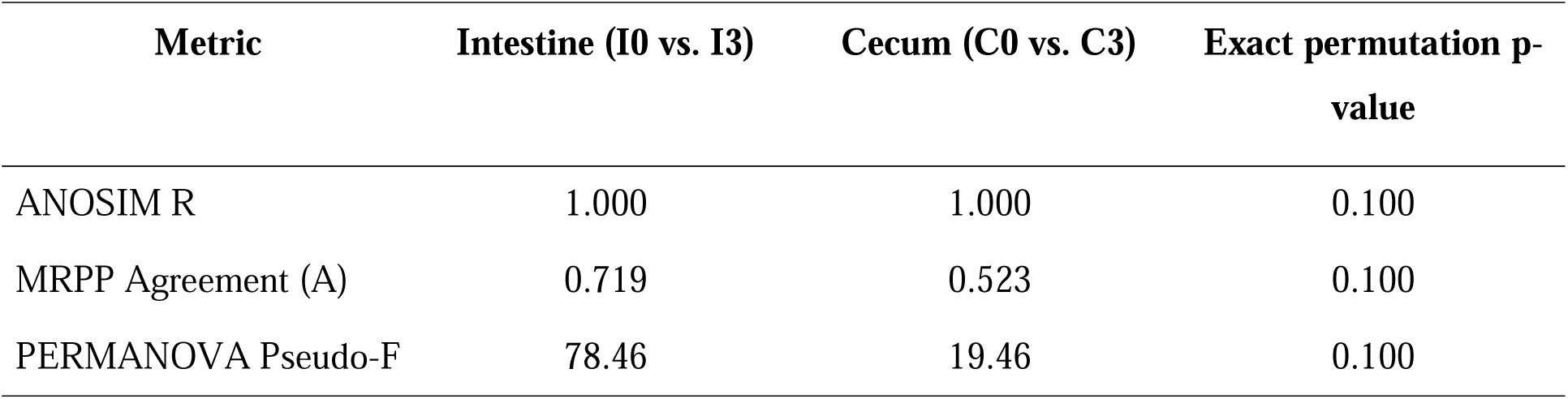
Statistical metrics for community separation (control vs. betaine + sodium bicarbonate) by intestinal segment

| **Metric** | **Intestine (I0 vs. I3)** | **Cecum (C0 vs. C3)** | **Exact permutation p-value** |
|---|---|---|---|
| ANOSIM R | 1.000 | 1.000 | 0.100 |
| MRPP Agreement (A) | 0.719 | 0.523 | 0.100 |
| PERMANOVA Pseudo-F | 78.46 | 19.46 | 0.100 |

A similar clustering pattern was observed using Bray–Curtis dissimilarity, computed independently for each intestinal site (Figure 4). In both tissue types, control and treatment groups formed completely segregated clusters along the first principal coordinate axis. PCoA1 accounted for 84.21% of the total dissimilarity in the cecum and 94.99% in the small intestine.

**Figure 4.**
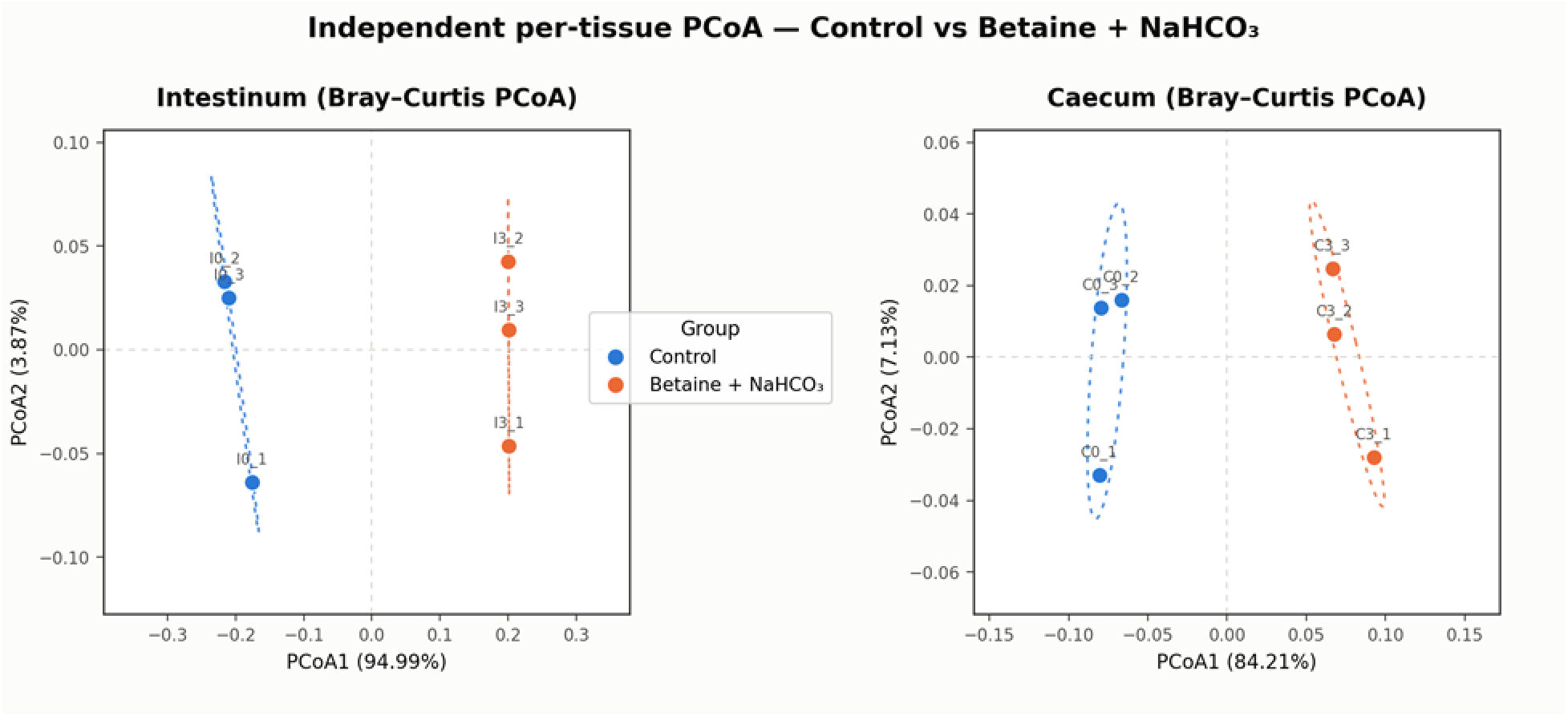
Site-specific PCoA based on Bray–Curtis dissimilarity computed independently for small intestinal samples (left) and cecal samples (right). Blue = control; Orange = betaine + sodium bicarbonate treatment.

#### Species-level microbial composition

Species-level relative abundances changed significantly between T0 and T3 in both the small intestine and cecum (Figure 5). A total of 9 species in the intestine and 5 species in the cecum showed statistically significant differences. In the intestine, *Aeriscardovia aeriphila* was the most enriched species, with its relative abundance increasing from 16% to 50% (*p* < 0.01). Conversely, *Lactobacillus delbrueckii* decreased from 46% to 32% (*p* < 0.01). Several species were effectively eliminated from the intestinal community in T3 relative to T0. These included *Enterococcus cecorum* (from 7.5% to undetectable; *p* < 0.01), *Streptococcus ferus* (from 4% to undetectable; *p* < 0.01), *Romboutsia sedimentorum* (from 4% to undetectable; *p* < 0.01), and *Streptococcus hyovaginalis* (from 3.5% to undetectable; p < 0.01). Additionally, *Romboutsia ilealis* declined significantly from 5% to 1% (p < 0.01). On the other hand, two species emerged or expanded significantly in the T3 group: *Corticicoccus populi*, which increased from undetectable to 5% (*p* < 0.01), and *Turicibacter sanguinis*, which increased modestly from 3.5% to 5.5% (*p* < 0.05).

**Figure 5.**
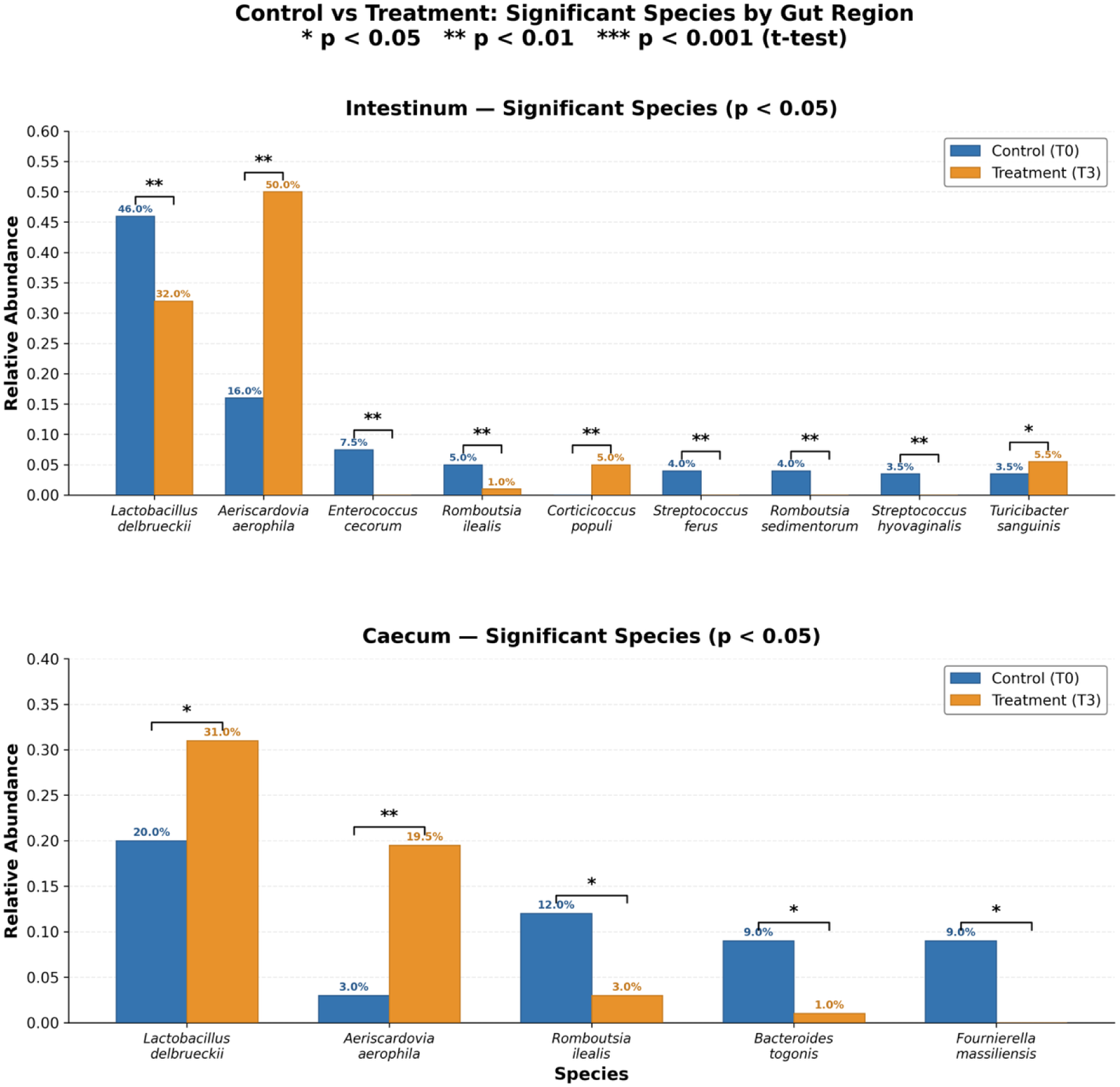
Relative abundance in bacterial species between control (T0) and treatment (T3) groups in the intestine (upper panel) and cecum (lower panel). Significance determined by unpaired Student’s *t*-test: * *P* < 0.05; ** *P* < 0.01.

In the cecum, supplementation with T3 significantly increased the proportion of two species: *Lactobacillus delbrueckii*, which rose from 20% to 31% (*p* < 0.05), and *Aeriscardovia aeriphila*, which went up by about 6.5-fold from 3% to 19.5% (*p* < 0.01). On the other hand, three species showed a significant decrease: *Romboutsia ilealis* fell from 12% to 3% (*p* < 0.05), *Bacteroides togonis* was almost eliminated, dropping from 9% to about 1% (*p* < 0.05), and *Fournierella massiliensis* decreased from 9% to undetectable levels (*p* < 0.05).

## Discussion

### Environmental conditions

The environmental conditions involved a constant thermal stress during the experiment; the average THI was 29.68, the weekly values ranging from 28.71 to 30.74 and daily temperatures frequently exceeding 30°C. As indicated by the standard THI classification, most of the recorded days fell into either the severe or very severe heat stress categories (Habeeb et al. 2018). These conditions provide a reliable foundation for evaluating nutritional strategies intended for maintaining physiological and productive performance in laying hens exposed to prolonged thermal stress. It is well known that heat stress in poultry results in a rise in body temperature, causes a decrease in feed intake, impairs nutrient utilization, alters the structure of the intestine and disturbs the gastrointestinal microbial community (Burkholder et al. 2008; Purswell et al. 2012; Bhadauria et al. 2014; Nawab et al. 2018; Ahmad et al. 2022). More recent evidence has demonstrated that heat stress also affects the microbial diversity and the community structure in the gastrointestinal tract of chickens (Ncho 2025).

The relatively high and persistent THI that was observed in this study is particularly important since the experiment was conducted in a tropical production environment and not as a result of a short-term experimental heat stress. It follows that the changes in rectal temperature, production performance, intestinal morphology, and microbial composition should all be understood in the light of chronic environmental exposure.

### Rectal temperature

The rectal temperature was lowered when the hens were given a dietary supplement of betaine plus sodium bicarbonate, with the most consistent decreases being observed at T3 and T4 in the later part of the experiment. During the 7-week assessment period, the average temperature at T3 was 41.20°C and that at T4 was 41.37°C, as compared with 41.57°C in the control group (T0). The fact that these groups showed a continued lowering of rectal temperature indicates that the combination of betaine and sodium bicarbonate assisted the hens in maintaining their body temperature under control during thermal stress.

The results support earlier studies which showed that betaine functions as an osmoprotective agent (Ratriyanto et al. 2024). It mainly acts as a compatible osmolyte and this helps to maintain cellular water and electrolyte balance under different environmental stresses (Ratriyanto and Mosenthin 2018; Abd El-Ghany and Babazadeh 2022). Experiments with laying poultry at high ambient temperatures showed a reduction in rectal temperature after they were given betaine in their diet (Ratriyanto et al. 2024). In particular, Al-Qaisi et al. (2023) reported a decrease in rectal temperature in heat-stressed laying hens receiving 2 g/kg betaine, along with improved egg production and feed efficiency. Similar effects have also been observed in broiler chickens, because the addition of betaine lessened the thermoregulatory strain and enhanced intestinal tissue integrity when the birds were exposed to heat (Shakeri et al. 2018).

Sodium bicarbonate is important in maintaining electrolyte levels and the acid-base balance when animals are exposed to heat, mainly because it supplies both bicarbonate and sodium. When poultry are under heat stress, they generally increase their respiratory rate to dissipate heat, a process that can disrupt the balance between carbon dioxide and bicarbonate in the body, thereby causing respiratory alkalosis. Sodium bicarbonate supplementation increases the dietary electrolyte balance without adding chloride, stimulates water intake, and supplies carbonate for eggshell formation, all of which may help birds cope with prolonged panting (Livingston et al. 2022). Consequently, dietary supplementation with electrolytes, including sodium bicarbonate, has been explored as a nutritional intervention to support physiological stability under elevated temperatures (Mushtaq et al. 2013; Livingston et al. 2022). Furthermore, empirical evidence indicates that sodium bicarbonate supplementation can affect thermoregulation and acid–base balance in heat-stressed poultry (Ahmad et al. 2008). These findings are consistent with the proposed physiological roles of sodium bicarbonate and betaine in supporting thermoregulation.

### Production performance

The supplements exerted significant effects on FI, HDP, EW, EM, and FCR. Treatment group T3 exhibited the highest HDP (89.93%) and the lowest FCR (1.85), despite a lower FI than the control group (*p* < 0.05). The lower feed conversion ratio in T3, together with its improved intestinal morphology, suggests more efficient nutrient utilization underlying the enhanced production response. The observed improvements in laying performance align with previous studies indicating a favorable influence of betaine under heat stress. Al-Qaisi et al. (2023) demonstrated that betaine supplementation enhanced HDP and EM in laying hens subjected to heat stress, while also reducing FCR in comparison to unsupplemented heat-stressed birds. Similarly, Liu et al. (2019) reported that dietary betaine enhanced nutrient utilization and performance in heat-stressed poultry. These positive effects are mainly due to betaine’s osmoprotective properties and its function as a methyl donor, which may assist in maintaining cellular homeostasis and nutrient efficiency under thermal stress (Ratriyanto and Mosenthin 2018; Ghasemi and Nari 2020). The better performance of T3 compared to T1 suggests that sodium bicarbonate contributed to the response, because both treatments contained the same betaine concentration; however, as no sodium bicarbonate-only treatment was included, an additive or synergistic interaction cannot be inferred.

The reduced FI across all groups given the supplements should be taken into account when evaluating FCR. Normally, under high ambient temperatures, FI decreases in laying hens to reduce metabolic heat production, thereby limiting the availability of nutrients for egg production (Ratriyanto and Mosenthin 2018). In this study, however, the T3 group kept the highest HDP even though the FI was less than that of the control group. This indicates an improved FCR when birds were supplemented with betaine together with sodium bicarbonate.

The higher EW observed in T4 as compared to T3 shows that the responses to each individual production trait were not similar under the different supplementation regimens: the treatment which resulted in the highest HDP was not necessarily the one that generated the greatest EW. It is possible that adding vitamin C to T4 caused the increased EW. Vitamin C has been widely studied as an antioxidant nutrient during heat stress, especially in cases where the bird’s own synthesis is unable to meet the increased physiological demands (El-Senousey et al. 2018; Van Hieu et al. 2022).

The value of EM, which is the result of the combined effect of HDP and EW, was greatest in T3 (54.83 g) and also in T4 (54.30 g), the two treatments being similar to each other even though they achieved this result via different production components: in T3, HDP was the highest, but EW was not the highest, while in T4, EW was the highest but HDP was lower than in T3. This indicates that the improvement in EM achieved through co-supplementation with sodium bicarbonate in T3 was mainly due to a higher laying rate, whereas in T4 the improvement in EM from co-supplementation with vitamin C was mainly due to larger egg size. Al-Qaisi et al. (2023) also reported a similar improvement in EM when betaine was supplemented under heat stress, attributing this increase mainly to an improved HDP rather than to EW, a finding which is in agreement with the pattern in this study for T3. However, the route to higher EM via EW observed in T4 has not been commonly reported in betaine-based diets, implying that vitamin C’s antioxidant effect may operate through a mechanism different from betaine’s osmoprotective and methyl-donor actions.

### Intestinal histomorphology

The most striking morphological differences between T0 and T3 were found in the duodenum and jejunum, which are the most important sites of digestion and absorption (Table 4). In the duodenum, T3 enhanced VH from 2,158 µm to 3,030 µm and VW from 335 µm to 761 µm, along with an approximately 3.2-fold rise in the absorptive surface area (from 2.26 to 7.26 mm²). In the jejunum, VH increased from 1,144 µm to 1,962 µm and VW from 187 µm to 493 µm, which led to a 4.5-fold increase in the absorptive surface area (from 0.67 to 3.03 mm²). In the ileum, however, there was no significant difference between treatments in terms of VH and CD, but both VW and the absorptive surface area were higher in T3 than in T0, showing that the ileum’s response to supplementation was less pronounced than in the duodenum and jejunum and that it was mainly due to villus widening rather than an increase in VH or CD.

These results are consistent with earlier studies that have shown that heat stress negatively affects intestinal structure, generally leading to a decrease in VH and an increase in CD. It has been demonstrated that prolonged exposure to heat reduces jejunal VH and the villus-to-crypt ratio in poultry, suggesting a disturbance of the mucosal structure (He et al. 2018). In contrast, supplementation with betaine has been linked to maintaining or improving intestinal structure under heat stress (Rahma et al. 2026). Liu et al. (2019) observed that betaine counteracted the heat stress-induced decreases in VH and the villus-to-crypt ratio in broilers. Shakeri et al. (2018) also reported that intestinal integrity and thermoregulation were improved as a result of betaine supplementation.

The observed response in this study aligns with the osmoprotective function of betaine, which can accumulate in intestinal tissues, thereby stabilizing epithelial cells under osmotic stress (Ratriyanto and Mosenthin 2018). Kettunen et al. (2001) showed that dietary betaine accumulates in intestinal tissue and affects intestinal epithelial structure. Recent investigations in aged laying hens have similarly linked dietary betaine to increases in VH and villus-to-crypt ratio, as well as alterations in the expression of intestinal barrier-related proteins (Du et al. 2025). Furthermore, the extra buffering capacity provided by sodium bicarbonate likely enhances this effect by keeping luminal pH at a level that reduces acid-induced erosion of the epithelium and promotes proliferative activity of crypt stem cells (Livingston et al. 2022).

The histomorphological effects were clearly specific to certain segments. In the duodenum and jejunum, there were changes in both VH and VW, whereas in the ileum, there was primarily an increase in VW. Since the absorptive surface area takes into account the dimensions of the villi, these alterations led to a considerably greater estimated mucosal surface area in T3. However, it is essential to consider the surface area as a morphometric estimate rather than as direct proof of increased nutrient absorption. Exercise caution when interpreting the rise in CD in the duodenum and jejunum. Although deeper crypts could suggest enhanced epithelial renewal, CD alone cannot confirm increased regenerative activity. Therefore, direct measurements such as Ki-67 expression, epithelial turnover rates, or crypt-cell proliferation are needed to support this mechanism (Marchini et al. 2011; Li et al. 2018; Mohammed 2019).

Previous studies including heat-stressed laying hens have not consistently shown that betaine restores intestinal morphology. For instance, Al-Qaisi et al. (2023) discovered that betaine had no effect on the intestinal morphology although it enhanced HDP. The more prominent morphological response observed in this study could be attributable to the additional inclusion of sodium bicarbonate.

### Intestinal microbiota composition

The results from the microbiota analysis showed significant variations according to intestinal site and dietary group. Alpha diversity values were lower in the intestine of the T3 group than in the T0 group, while the cecal Chao1, Shannon, and Simpson indices were mostly similar across treatments. This pattern, which varies by segment, is in agreement with the well-documented ecological differences between the proximal small intestine and the cecum in chickens, since these organs harbor different microbial communities that correspond to differences in the availability of substrate, transit time, oxygen tension and the physicochemical conditions (Khan et al. 2020; Rychlik 2020). It is also necessary to stress that a decrease in alpha diversity should not be taken automatically as an indication of worse gut health. Since diversity is an ecological characteristic rather than a direct indicator of host health or of the functional capacity of the microorganisms, a microbial community with less taxonomic diversity can still maintain, or even improve, certain functional abilities. This point is especially relevant in this case, since the intestinal microbial response was characterized by great changes in a small number of particular taxa rather than by a general loss of community structure.

The analysis of beta diversity confirmed a similar pattern specific to segments. In both the intestine and the cecum, the T0 and T3 groups formed distinct clusters (with an ANOSIM R value of 1.000 for both sites), although the degree of separation varied by site. The PERMANOVA pseudo-F value was more than four times greater in the small intestine than in the cecum, and complete clustering in the intestine was already apparent along the first, main coordinate axis, whereas separation of the cecal populations had to be looked at along a minor axis. This shows that the supplementation with betaine and sodium bicarbonate had a much stronger effect on the intestinal microbiota than on the cecal microbiota, which is in agreement with the small intestine being the main site of absorption of nutrients and osmolytes and thus the site most directly affected by the osmotic and pH-modulating effects of the supplements (Ratriyanto and Prastowo 2019).

At the species level, *Aeriscardovia aeriphila* gained the most from the supplementation, showing a considerable increase in relative abundance in both the intestine and the cecum. As a member of the family *Bifidobacteriaceae*, *A. aeriphila* ferments carbohydrates via the bifid shunt, producing acetate and lactate as its main end products; earlier studies have shown that giving this species orally improves the growth performance, antioxidant capacity, and immune function of broilers by promoting the proliferation of *Firmicutes* (Farooq et al. 2023). The osmoprotective effect of betaine combined with the pH-buffering property of sodium bicarbonate most likely created conditions favorable for the proliferation of the probiotic species by ensuring that the substrate remained available while preventing pH-sensitive bacteria from growing.

The observation that *Lactobacillus delbrueckii* exhibited a site-specific response—decreasing in the intestine but increasing in the cecum—is among the more ecologically informative. The fact that *L. delbrueckii* is a homofermentative lactic acid bacterium well adapted to low-pH environments (Plaza-Diaz et al. 2019; Wang et al. 2021) makes a pH-buffering explanation for its cecal increase unlikely; the increase may instead reflect altered substrate flow to the ceca. In the intestine, however, the vigorous proliferation of *A. aeriphila* may have resulted in competitive exclusion by using up the available substrates and causing niche overlap, in line with the ecological principle that closely related species occupying the same functional niche cannot coexist indefinitely when competing for resources (Segura Munoz et al. 2022). In engineered anaerobic digesters, co-dominance of *Aeriscardovia* and *Lactobacillus* has been associated with acetate and lactate production; whether this applies to the avian gut remains to be determined (Moestedt et al. 2020), which shows that the intestinal site was an important factor in determining the microbial response even among taxa within the same saccharolytic functional group (Khan et al. 2020; Rychlik 2020).

The near-elimination of *Bacteroides togonis* and the decline of *Romboutsia ilealis* in the cecum may indicate a reduction in taxa with proteolytic potential. *Bacteroides* species have substantial proteolytic potential, and a serine protease from *Bacteroides fragilis* has been shown to cleave and activate host protease-activated receptor 2, thereby affecting epithelial integrity (Obiso et al. 1995; Lakemeyer et al. 2025). Their decline may therefore reflect a protective microbial adaptation that reduces host exposure to protease-mediated epithelial disruption. Similarly, *Romboutsia* is a genus associated with amino acid degradation and bile acid metabolism (He et al. 2025), and declined in parallel with the broader shift away from protein putrefaction toward carbohydrate fermentation.

A notable finding was the complete disappearance of several intestinal species, including *Enterococcus cecorum* and two *Streptococcus* species. *E. cecorum* normally resides in the avian gut as a commensal, but pathogenic strains have emerged as significant causes of septicemia and skeletal issues, including vertebral osteomyelitis and femoral head necrosis (Borst et al. 2017; Jung et al. 2018). Thus, its elimination during T3 supplementation suggests a potential pathogen-suppressive effect of the dietary intervention. The concurrent loss of both Streptococcus species is consistent with the well-established antimicrobial mechanism of lactic acid bacteria, in which organic acid production lowers luminal pH and competitively excludes acid-sensitive taxa (Reuben et al. 2019; Tang et al. 2023). Additionally, the exclusive presence of *Corticicoccus populi* in the T3 intestine illustrates the opportunistic niche exploitation commonly observed after resident proteolytic and streptococcal populations are displaced (Khan et al. 2020).

Taken together, the species-level changes suggest a shift from taxa with proteolytic potential toward saccharolytic taxa. The enrichment of *Aeriscardovia* and *L. delbrueckii*, together with the suppression of *R. ilealis, B. togonis*, and several species of *Streptococcus*, is consistent with a shift toward saccharolytic rather than proteolytic fermentation, possibly mediated by the acidifying and pH-lowering effects of lactic and acetic acid produced by *Lactobacillus* and *Aeriscardovia* (Tang et al. 2023; Boven et al. 2025). This putative benefit, however, came at the cost of reduced taxonomic diversity in the intestine, illustrating a trade-off between ecological simplification and functional specialization that should be considered when optimizing future betaine- and sodium bicarbonate-based supplementation strategies.

## Conclusions

Under chronic tropical heat stress, betaine combined with sodium bicarbonate (T3) produced the most favorable production response, lowering rectal temperature relative to the control and achieving the highest hen-day egg production (89.93%) and egg mass (54.83 g) and the best feed conversion ratio (1.85). Compared with the control, T3 improved intestinal mucosal architecture, with larger villus dimensions and greater calculated absorptive surface area in the duodenum and jejunum, and increased villus width and absorptive surface area in the ileum, although nutrient absorption was not measured directly. The combination also restructured the gut microbiota in a site-dependent manner. Intestinal alpha diversity decreased and community separation was stronger than in the cecum, where diversity was largely maintained. Enrichment of *Aeriscardovia aeriphila* and reductions in *Romboutsia ilealis*, *Bacteroides togonis*, *Enterococcus cecorum*, and several *Streptococcus* species suggest a shift in the balance of saccharolytic and proteolytic-associated taxa. Overall, betaine plus sodium bicarbonate is a promising strategy for sustaining laying performance and intestinal morphology under heat stress. Future factorial studies should optimize dose and ratio and directly assess microbial metabolites, nutrient digestibility, and functional capacity.

## Acknowledgements

Funding for this work was provided by Universitas Sebelas Maret through the “*Penelitian Fundamental B (PFB-UNS)*” scheme, in accordance with Research Assignment Agreement No. 460/UN27.22/PT.01.03/2026.

## Author contributions

ZAW: Conceptualization, Investigation, Formal analysis, Writing – original draft; AR: Conceptualization, Supervision, Funding acquisition, Writing – review & editing; EAS: Methodology, Writing – review & editing; SP: Formal analysis, Writing – review & editing; LAP: Investigation, Data curation.

## Data availability statement

The raw 16S rRNA gene amplicon sequencing data and other datasets generated during the current study are not publicly available because they form part of ongoing research, but are available from the corresponding author upon reasonable request.

## References

1. Abbas G, Mahmood S, Haq A ul, Nawaz H. 2022. Effect of Dietary Inclusion of Sodium Bicarbonate on Production Performance o Caged Layers during Summer. Pak J Zool. 54(2):751–759. 10.17582/journal.pjz/20190705200740

2. Abd El-Ghany WA, Babazadeh D. 2022. Betaine: A Potential Nutritional Metabolite in the Poultry Industry. Animals. 12(19):2624. 10.3390/ani12192624

3. Abdullah BJ, Hamza MAA, Rasool MH. 2025. Effect Adding of Sodium Bicarbonate in Broiler Water On Performance and Immune Response. J Neonatal Surg. 14:1304–1309. 10.52783/jns.v14.1955

4. Ahmad R, Yu YH, Hsiao FSH, Su CH, Liu HC, Tobin I, Zhang G, Cheng YH. 2022. Influence of heat stress on poultry growth performance, intestinal inflammation, and immune function and potential mitigation by probiotics. Animals. 12(17):2297. 10.3390/ani12172297

5. Ahmad T, Khalid T, Mushtaq T, Mirza MA, Nadeem A, Babar ME, Ahmad G. 2008. Effect of potassium chloride supplementation in drinking water on broiler performance under heat stress conditions. Poult Sci. 87(7):1276–1280. 10.3382/ps.2007-00299

6. Al Wahid Z, Pradista LA, Prastowo S, Ratriyanto A. 2024. Intestinal bacterial composition in laying pullet reared in tropical climate received betaine supplementation. IOP Conf Ser Earth Environ Sci. 1362(1). 10.1088/1755-1315/1362/1/012046

7. Al-Qaisi M, Abdelqader A, Abuajamieh M, Abedal-Majed MA, Al-Fataftah ARA. 2023. Impacts of dietary betaine on rectal temperature, laying performance, metabolism, intestinal morphology, and follicular development in heat-exposed laying hens. J Therm Biol. 117:103714. 10.1016/j.jtherbio.2023.103714

8. Attia YA, Abd El-Hamid AEHE, Abedalla AA, Berika MA, Al-Harthi MA, Kucuk O, Sahin K, Abou-Shehema BM, Diarra SS, Tabuaciri P, et al. 2021. Laying performance, digestibility and plasma hormones in laying hens exposed to chronic heat stress as affected by betaine, vitamin C, and/or vitamin E supplementation. J Therm Biol. 5(1):576–595. 10.1186/s40064-016-3304-0

9. Bhadauria P, Kataria JM, Majumdar S, Bhanja SK, Kolluri G. 2014. Impact of Hot Climate on Poultry Production System-A Review Thermoregulatory Mechanism of Poultry. J Poult Sci Technol. 2(4):56–63.

10. Borst LB, Suyemoto MM, Sarsour AH, Harris MC, Martin MP, Strickland JD, Oviedo EO, Barnes HJ. 2017. Pathogenesis of Enterococcal Spondylitis Caused by Enterococcus cecorum in Broiler Chickens. Vet Pathol. 54(1):61–73. 10.1177/0300985816658098

11. Boven L, Akkerman R, de Vos P. 2025. Sustainable diets with plant-based proteins require considerations for prevention of proteolytic fermentation. Crit Rev Food Sci Nutr. 65(14):2829–2839. 10.1080/10408398.2024.2352523

12. Burkholder KM, Thompson KL, Einstein ME, Applegate TJ, Patterson JA. 2008. Influence of stressors on normal intestinal microbiota, intestinal morphology, and susceptibility to Salmonella Enteritidis colonization in broilers. Poult Sci. 87(9):1734–1741. 10.3382/ps.2008-00107

13. Chen R, Yang M, Song YD, Wang RX, Wen C, Liu Q, Zhou YM, Zhuang S. 2022. Effect of anhydrous betaine and hydrochloride betaine on growth performance, meat quality, postmortem glycolysis, and antioxidant capacity of broilers. Poult Sci. 101(4):101687. 10.1016/j.psj.2021.101687

14. Du X, Pian H, Zhao D, Zhang Y, Wu X, He J, Chen L, Liu F, Yu D. 2025. Enhancing gut-ovary health in aged laying hens: the impact of dietary betaine supplementation. Poult Sci. 104(4):104894. 10.1016/j.psj.2025.104894

15. El-Senousey HK, Chen B, Wang JY, Atta AM, Mohamed FR, Nie QH. 2018. Effects of dietary Vitamin C, Vitamin E, and alpha-lipoic acid supplementation on the antioxidant defense system and immune-related gene expression in broilers exposed to oxidative stress by dexamethasone. Poult Sci. 97(1):30–38. 10.3382/ps/pex298

16. Farag MR, Alagawany M. 2018. Physiological alterations of poultry to the high environmental temperature. J Therm Biol. 76:101–106. 10.1016/j.jtherbio.2018.07.012

17. Farooq MZ, Wang X, Yan X. 2023. Effects of Aeriscardovia aeriphila on growth performance, antioxidant functions, immune responses, and gut microbiota in broiler chickens. J Zhejiang Univ Sci B. 24(11):1014–1026. 10.1631/jzus.B2200621

18. Ghasemi HA, Nari N. 2020. Effect of supplementary betaine on growth performance, blood biochemical profile, and immune response in heat-stressed broilers fed different dietary protein levels. J Appl Poult Res. 29(2):301–313. 10.1016/j.japr.2019.11.004

19. Habeeb AA, Gad AE, Atta MA. 2018. Temperature-Humidity Indices as Indicators to Heat Stress of Climatic Conditions with Relation to Production and Reproduction of Farm Animals. Int J Biotechnol Recent Adv. 1(1):35–50. 10.18689/ijbr-1000107

20. Hassan MSH, El-Sayed OA, Namera MMM. 2011. Effect of dietary sodium bicarbonate and potassium chloride supplementation on acid-base balance, plasma electrolytes and aldosterone hormone of Golden Montazah hens under hot climate condition. Egypt Poult Sci J. 31(2):285–303.

21. He X, Lu Z, Ma B, Zhang L, Li J, Jiang Y, Zhou G, Gao F. 2018. Effects of chronic heat exposure on growth performance, intestinal epithelial histology, appetite-related hormones and genes expression in broilers. J Sci Food Agric. 98(12):4471–4478. 10.1002/jsfa.8971

22. He Z, Guo M, Zhang X, Wang S, Liu T, Lin Y, Ouyang Q, Hu S, He H, Li L, et al. 2025. 16S rRNA and transcriptome analysis revealed the regulatory mechanism of Romboutsia lituseburensis on serum immunoglobulin levels in geese. Poult Sci. 104(5):105018. 10.1016/j.psj.2025.105018

23. Jung A, Chen LR, Suyemoto MM, Barnes HJ, Borst LB. 2018. A Review of Enterococcus cecorum Infection in Poultry. Avian Dis. 62(3):261–271. 10.1637/11825-030618-Review.1

24. Kaya I, Karademir B, Ucar O. 2004. The effects of diet supplemented with sodium bicarbonate upon blood pH, blood gases and eggshell quality in laying geese. Vet Med (Praha). 49(6):201–206. 10.17221/5696-VETMED

25. Kettunen H, Tiihonen K, Peuranen S, Saarinen MT, Remus JC. 2001. Dietary betaine accumulates in the liver and intestinal tissue and stabilizes the intestinal epithelial structure in healthy and coccidia-infected broiler chicks. Comp Biochem Physiol A Mol Integr Physiol. 130(4):759–769. 10.1016/S1095-6433(01)00410-X

26. Khan S, Moore RJ, Stanley D, Chousalkar KK. 2020. The gut microbiota of laying hens and its manipulation with prebiotics and probiotics to enhance gut health and food safety. Appl Environ Microbiol. 86(13):e00600–20. 10.1128/AEM.00600-20

27. Kim DY, Kim RH, Kim HW, Lee JH, Kil DY. 2026. Effect of dietary glycine and betaine on productive performance, liver health, intestinal characteristics, and stress response in aged laying hens under heat stress conditions. Anim Biosci. 39(5). 10.5713/ab.250618

28. Lakemeyer M, Latorre R, Blazkova K, Jensen D, Wood HM, Shakil N, Thomas SC, Saxena D, Malpuri Y, Poolman D, et al. 2025. Identification of a secreted protease from Bacteroides fragilis that induces intestinal pain and inflammation by cleavage of PAR2. 10.1101/2025.01.15.633241

29. Li J, Li J, Zhang SY, Li RX, Lin X, Mi YL, Zhang CQ. 2018. Culture and characterization of chicken small intestinal crypts. Poult Sci. 97(5):1536–1543. 10.3382/ps/pey010

30. Liu W, Yuan Y, Sun C, Balasubramanian B, Zhao Z, An L. 2019. Effects of dietary betaine on growth performance, digestive function, carcass traits, and meat quality in indigenous yellow-feathered broilers under long-term heat stress. Animals. 9(8):506. 10.3390/ani9080506

31. Livingston ML, Pokoo-Aikins A, Frost T, Laprade L, Hoang V, Nogal B, Phillips C, Cowieson AJ. 2022. Effect of Heat Stress, Dietary Electrolytes, and Vitamins E and C on Growth Performance and Blood Biochemistry of the Broiler Chicken. Front Anim Sci. 3:807267. 10.3389/fanim.2022.807267

32. Marchini CFP, Silva PL, Nascimento MRBM, Beletti ME, Silva NM, Guimaraes EC. 2011. Body Weight, Intestinal Morphometry and Cell Proliferation of Broiler Chickens Submitted to Cyclic Heat Stress. Int J Poult Sci. 10(6):455–460. 10.3923/ijps.2011.455.460

33. Martínez Y, Almendares CI, Hernández CJ, Avellaneda MC, Urquía AM, Valdivié M. 2021. Effect of acetic acid and sodium bicarbonate supplemented to drinking water on water quality, growth performance, organ weights, cecal traits and hematological parameters of young broilers. Animals. 11(7):1865. 10.3390/ani11071865

34. Moestedt J, Müller B, Nagavara Nagaraj Y, Schnürer A. 2020. Acetate and Lactate Production During Two-Stage Anaerobic Digestion of Food Waste Driven by Lactobacillus and Aeriscardovia. Front Energy Res. 8:105. 10.3389/fenrg.2020.00105

35. Mohammed AA. 2019. Quantitative assessment of Ki67 expression in correlation with various breast cancer characteristics and survival rate; cross sectional study. Ann Med Surg. 48:129–134. 10.1016/j.amsu.2019.11.005

36. Mushtaq MMH, Pasha TN, Mushtaq T, Parvin R. 2013. Electrolytes, dietary electrolyte balance and salts in broilers: An updated review on growth performance, water intake and litter quality. Worlds Poult Sci J. 69(4):789–802. 10.1017/S0043933913000810

37. Nawab A, Ibtisham F, Li G, Kieser B, Wu J, Liu W, Zhao Y, Nawab Y, Li K, Xiao M, An L. 2018. Heat stress in poultry production: Mitigation strategies to overcome the future challenges facing the global poultry industry. J Therm Biol. 78:131–139. 10.1016/j.jtherbio.2018.08.010

38. Ncho CM. 2025. Heat stress and the chicken gastrointestinal microbiota: a systematic review. J Anim Sci Biotechnol. 16(1):85. 10.1186/s40104-025-01225-6

39. Obiso RJ, Lyerly DM, Van Tassell RL, Wilkins TD. 1995. Proteolytic activity of the Bacteroides fragilis enterotoxin causes fluid secretion and intestinal damage in vivo. Infect Immun. 63(10):3820–3826. 10.1128/iai.63.10.3820-3826.1995

40. Onagbesan OM, Uyanga VA, Oso O, Tona K, Oke OE. 2023. Alleviating heat stress effects in poultry: updates on methods and mechanisms of actions. Front Vet Sci. 10:1255520. 10.3389/fvets.2023.1255520

41. Plaza-Diaz J, Ruiz-Ojeda FJ, Gil-Campos M, Gil A. 2019. Mechanisms of Action of Probiotics. Adv Nutr. 10:S49–S66. 10.1093/advances/nmy063

42. Prakatur I, Miskulin M, Pavic M, Marjanovic K, Blazicevic V, Miskulin I, Domacinovic M. 2019. Intestinal Morphology in Broiler Chickens Supplemented with Propolis and Bee Pollen. Animals. 9(6):301. 10.3390/ani9060301

43. Purswell JL, Dozier WA, Olanrewaju HA, Davis JD, Xin H, Gates RS. 2012. Effect of temperature-humidity index on live performance in broiler chickens grown from 49 to 63 days of age. :41–49. 10.13031/2013.41619

44. Rahma SA, Ratriyanto A, Prastowo S, Sayogo AD. 2026. Growth performance, intestinal morphometrics, and nutrient digestibility of betaine-supplemented Kedu chickens reared at different altitudes. Open Vet J. 16(6):3678–3689. 10.5455/OVJ.2026.v16.i6.39

45. Ratriyanto A, Mosenthin R. 2018. Osmoregulatory function of betaine in alleviating heat stress in poultry. J Anim Physiol Anim Nutr (Berl). 102(6):1634–1650. 10.1111/jpn.12990

46. Ratriyanto A, Prastowo S, Sofyan A, Widyas N, Pradista LA. 2024. Osmoprotective function of betaine maintains the physiological parameters of quail raised in tropical environments. IOP Conf Ser Earth Environ Sci. 1360(1):012034. 10.1088/1755-1315/1360/1/012034

47. Ratriyanto A, Prastowo S. 2019. Floor space and betaine supplementation alter the nutrient digestibility and performance of Japanese quail in a tropical environment. J Therm Biol. 83:80–86. 10.1016/j.jtherbio.2019.05.008

48. Reuben RC, Roy PC, Sarkar SL, Alam RU, Jahid IK. 2019. Isolation, characterization, and assessment of lactic acid bacteria toward their selection as poultry probiotics. BMC Microbiol. 19(1). 10.1186/s12866-019-1626-0

49. Rychlik I. 2020. Composition and function of chicken gut microbiota. Animals. 10(1):103. 10.3390/ani10010103

50. Salem HM, Alqhtani AH, Swelum AA, Babalghith AO, Melebary SJ, Soliman SM, Khafaga AF, Selim S, El-Saadony MT, El-Tarabily KA, Abd El-Hack ME. 2022. Heat stress in poultry with particular reference to the role of probiotics in its amelioration: An updated review. J Therm Biol. 108:103302. 10.1016/j.jtherbio.2022.103302

51. Santos HB, Vieira DA, Souza LP, Santos AL, Santos FR, Araujo Neto FR. 2019. Application of non-linear mixed models for modelling the quail growth curve for meat and laying. J Agric Sci. 156(10):1216–1221. 10.1017/S0021859619000169

52. Segura Munoz RR, Mantz S, Martínez I, Li F, Schmaltz RJ, Pudlo NA, Urs K, Martens EC, Walter J, Ramer-Tait AE. 2022. Experimental evaluation of ecological principles to understand and modulate the outcome of bacterial strain competition in gut microbiomes. ISME J. 16(6):1594–1604. 10.1038/s41396-022-01208-9

53. Shakeri M, Cottrell JJ, Wilkinson S, Ringuet M, Furness JB, Dunshea FR. 2018. Betaine and antioxidants improve growth performance, breast muscle development and ameliorate thermoregulatory responses to cyclic heat exposure in broiler chickens. Animals. 8(10):162. 10.3390/ani8100162

54. Tang H, Huang W, Yao YF. 2023. The metabolites of lactic acid bacteria: classification, biosynthesis and modulation of gut microbiota. Microb Cell. 10(3):49–62. 10.15698/mic2023.03.792

55. Van Hieu T, Guntoro B, Qui NH, Quyen NTK, Al Hafiz FA. 2022. The application of ascorbic acid as a therapeutic feed additive to boost immunity and antioxidant activity of poultry in heat stress environment. Vet World. 15(3):685–693. 10.14202/vetworld.2022.685-693

56. Wang XL, Liu ZY, Li YH, Yang LY, Yin J, He JH, Hou DX, Liu YL, Huang XG. 2021. Effects of Dietary Supplementation of Lactobacillus delbrueckii on Gut Microbiome and Intestinal Morphology in Weaned Piglets. Front Vet Sci. 8:692389. 10.3389/fvets.2021.692389

57. Whitehead CC, Keller T. 2003. An update on ascorbic acid in poultry. Worlds Poult Sci J. 59(2):161–184. 10.1079/WPS20030010

58. Yörük MA, Gül M, Hayirli A, Karaoglu M. 2004. Laying performance and egg quality of hens supplemented with sodium bicarbonate during the late laying period. Int J Poult Sci. 3(4):272–278. 10.3923/ijps.2004.272.278

59. Zulovich JM, DeShazer JA. 1990. Estimating egg production declines at high environmental temperatures and humidities. ASAE Paper No. 90-4021. St. Joseph (MI): American Society of Agricultural Engineers.

